# Extending Janus lectins architecture: characterization and application to protocells

**DOI:** 10.1101/2022.08.15.503968

**Authors:** Simona Notova, Lina Siukstaite, Francesca Rosato, Federica Vena, Aymeric Audfray, Nicolai Bovin, Ludovic Landemarre, Winfried Römer, Anne Imberty

## Abstract

Synthetic biology is a rapidly growing field with applications in biotechnology and biomedicine. Through various approaches, remarkable achievements, such as cell and tissue engineering, have been already accomplished. In synthetic glycobiology, the engineering of glycan binding proteins is being developed for producing tools with precise topology and specificity. We developed the concept of chimeric lectins, i.e., Janus lectin, with increased valency, and additional specificity. The novel engineered lectin, assembled as a fusion protein between the β-propeller domain from *Ralstonia solanacearum* and the β-trefoil domain from fungus *Marasmius oreades*, is specific for fucose and α-galactose and its unique protein architecture allows to bind these ligands simultaneously. The protein activity was tested with glycosylated giant unilamellar vesicles, resulting in the formation of proto-tissue-like structures through cross-linking of such protocells. The synthetic protein binds to H1299 lung epithelial cancer cells by its two domains. The biophysical properties of this new construct were compared with the two already existing Janus lectins, RSL-CBM40 and RSL-CBM77Rf. Denaturation profiles of the proteins indicate that the fold of each has a significant role in protein stability and should be considered during protein engineering.

## Introduction

Due to their ability to crosslink various cells, such are red blood cells, lectins are originally known as agglutinins. They are generally multivalent and their selective interaction with glycoconjugates found many applications in the field of biotechnology and biomedicine as glycan-profiling tools (Sharon and Lis 2004). Therefore, fine-tuning the specificity or the valency of lectins is a promising approach for obtaining novel tools. Synthetic biology strategies, such as the engineering of protein architecture, bring novel possibilities for lectin applications (Fettis et al. 2019; Irumagawa et al. 2022; Oh et al. 2022; Ramberg et al. 2021; Ross et al. 2019). Lectin engineering is a novel domain of synthetic glycobiology (Terada et al. 2016; Ward et al. 2021) and it can be performed at different levels, from glycan specificity to supramolecular architecture.

The trimeric oligomerization of *Ralstonia solanacearum* lectin (RSL) in a β-propeller shape (Fig. 1A) was recently used as a scaffold for building Janus lectins (Notova et al. 2022b; Ribeiro et al. 2018) with two faces and different specificities. The two Janus lectins, RSL-CBM40 and RSL-CBM77Rf, were obtained by fusion of the monomeric RSL with two carbohydrate-binding modules (CBMs) with different specificity but rather similar shape, consisting of a single β-sandwich domain. This resulted in the creation of synthetic bispecific chimeras able to establish the interaction with fucose on one side and sialic acid (RSL-CBM40) or homogalacturonan (RSL-CBM77_Rf_), respectively, on the other side.

**Figure 1:**
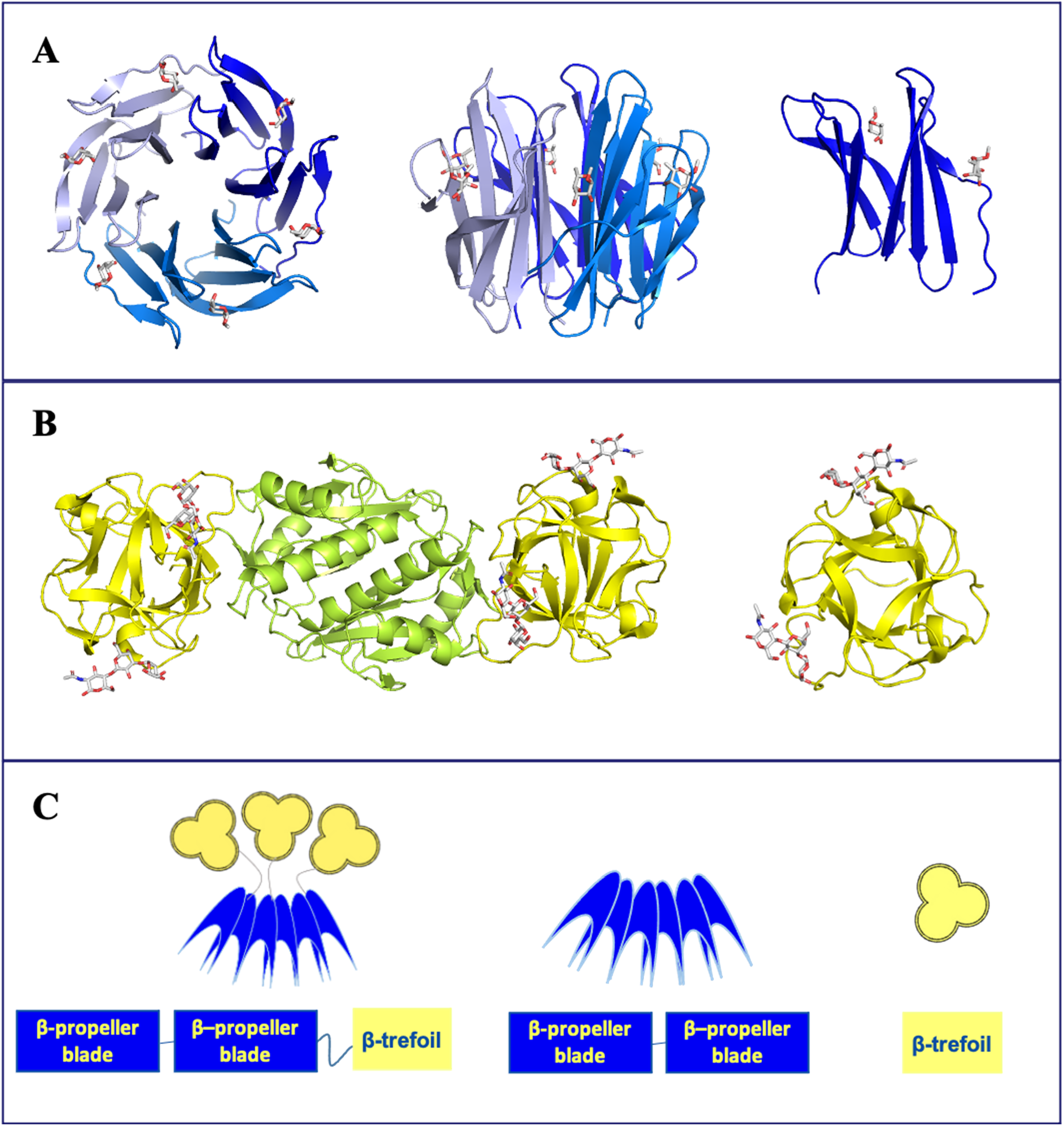
Schematic representations of Janus lectin RSL-MOA and crystal structures of its individual protein components. A) Left and middle: Top and side view of the crystal structure of trimeric RSL (blue carton) in complex with α-methyl fucoside (sticks) with in total of six fucose binding sites. Right: structure of RSL monomer (PDB code 2BT9). B) Left: Cartoon representation of the crystal structure of dimeric MOA, β-trefoil domain (yellow) in complex with Galα1-3Galβ1-4GlcNAc and dimerization domain (green). Right: β-trefoil domain of MOA with different orientations (PDB code 2IHO). C) Schematic representation of Janus lectin RSL-MOA, RSL, MOAβT (β-trefoil domain of MOA), and peptide domains.

Other protein domains with different specificity, but also topology diverging from the β-sandwich CBM, could be considered for building novel Janus lectins from the trimeric RSL scaffold. β-trefoils are robust domains that contain an internal tandem forming a three-lobed architecture with almost no secondary structures (Murzin et al. 1992). They display a large variety of functions, and many show carbohydrate-binding activity. They are considered as lectins due to their multivalence, but also as CBMs (e.g., CBM13) since they sometimes occur as domains associated with glyco-active enzymes or toxins (Notova et al. 2020; Notova et al. 2022a). While their shape is well conserved, their sequences do not show strong similarity, except for the presence of hydrophobic residues in the core and a QXW repeat found in most sequences (Hazes 1996). A large number of crystal structures of β-trefoil lectins are available (212 from 63 different proteins), and based on their sequences they have been classified into 12 classes in the UniLectin3D database (Bonnardel et al. 2019). Datamining of genomes based on the lobe sequence signature of each class resulted in the prediction of thousands of putative β-trefoil lectins spanning all kingdoms of life (Notova et al. 2022a). The threefold symmetry results in the presence of three carbohydrate-binding sites referred to as α, β, and γ although variations in amino acid sequences sometimes result in only one or two active binding sites.

Among the known structures of β-trefoil lectin, the agglutinin from the fairy ring mushroom *Marasmius oreades* (MOA) presents an interesting strict specificity for αGal containing oligosaccharides. The lectin rose attention already in the second half of the 20^th^ century due to its high selectivity and affinity to blood group B oligosaccharide whereas very poor binding was detected to blood group A or H oligosaccharides (Estola and Elo 1952; Winter et al. 2002).

MOA is specific for oligosaccharides with terminal non-reducing α1-3 linked galactose, while galactosides linked in α1-2, α1-4, and α1-6 showed no binding (Winter et al. 2002). Due to its preference for the αGal1-3Gal disaccharide, MOA binds efficiently to blood group B (Galα1-3[Fucα1-2]Gal) but also to linear oligosaccharides such as Galα1-3Galβ1-4Glc on glycosphingolipid isoGb3 and Galα1-3Galβ1-4GlcNAc (Galili epitope) on glycoproteins (Macher and Galili 2008). This latter epitope is present on the cell surface of most mammals, but not in humans, apes, and Old World monkeys due to the deactivation of the α3-galactosyltranferase during evolution (Galili et al. 1988). Humans possess preformed antibodies directed against Galα1-3Gal, which are responsible for hyperacute rejection of animal (mainly porcine) organs in xenotransplantation attempts (Cooper et al. 1994). They are also responsible for a strong immune response against biodrugs such as therapeutical antibodies if produced in rodent cell culture with active α1-3galactosyltranferase (Chung et al. 2008). Highly selective lectins are therefore needed for the identification of such epitopes (Kruger et al. 2002; Tateno and Goldstein 2004; Winter et al. 2002).

The crystal structure of MOA revealed that the lectin assembles as a homodimer with the monomer composed of two distinct domains, adopting β-trefoil fold at the N-terminus and α/β fold at the C-terminus (Fig. 1B). The C-terminal domain serves as a dimerization interface and retains a proteolytic function (Cordara et al. 2016; Juillot et al. 2016). The lectin β-trefoil domain has three-fold symmetry, and the conserved motif (Gln-X-Trp)3 is involved in the hydrophobic core of the structure. Co-crystals of MOA with Galα1-3Galβ1-4GlcNAc revealed that each binding site has different ligand occupancy, emphasizing the fact that slight differences in amino acids might affect the binding (Fig. 1B) (Grahn et al. 2007).

In the present study, we designed a Janus lectin as a fusion protein of monomeric RSL at the N-terminus and the MOA β-trefoil domain at the C-terminus (Fig. 1C). We produced, and characterized the Janus lectin RSL-MOA with double specificity toward fucosylated and α-galactosylated glycans. The ability to bind these epitopes was tested with H1299 lung epithelial cancer cells and giant unilamellar vesicles. We also compared the biophysical behavior of this novel Janus-lectin with RSL-CBM40 (Ribeiro et al. 2018) and RSL-CBM77_Rf_ (Notova et al. 2022b). Additionally, we have engineered the β-trefoil domain of MOA (MOAβT) and compared its activity with RSL-MOA.

## Results

### Design and production of the β-trefoil domain of MOA (MOAβT)

MOA occurs naturally as a dimer, assembled by a strong association between the two C-terminal domains. Monomeric MOA composed only of the β-trefoil domain would be of interest for biotechnology applications. The gene of the MOA β-trefoil domain was defined as the 156 N-terminal AAs of MOA full sequence and was amplified by PCR and subcloned into the vector pET TEV as a fusion with an N-terminal His-tag sequence. The resulting protein was named MOAβT and was recombinantly produced in soluble form in the bacterium *E. coli* BL21(DE3). The protein was purified by immobilized metal ion chromatography followed by His-tag cleavage by TEV (tobacco etch virus) protease. The resulting protein has an estimated molecular weight of 17.2 kDa (Supplementary Fig. 1A). TEV cleavage followed by additional immobilized metal ion chromatography reduced the sample contaminants and the protein purity was verified by 12% SDS PAGE electrophoresis (Supplementary Fig. 1A).

### Design and production of Janus lectin RSL-MOA

Janus lectin RSL-MOA was designed as a gene fusion of monomeric RSL (N-terminus) and β-trefoil domain of lectin MOA (C-terminus). The fusion of the two protein domains by the eight AAs linker PNGELLSS results in the protein sequence of 255 amino acids. The gene sequence was optimized for bacterial expression, and the synthesized gene was subcloned into the plasmid pET25b+. The protein was produced in soluble form in the bacterium *E. coli* KRX. The protein was subsequently purified by affinity chromatography on an agarose-mannose column due to the interaction between the RSL domain and mannose residues. Protein analysis by 12% SDS PAGE electrophoresis showed that Janus lectin RSL-MOA has an apparent size of 83 kDa, which corresponds to a trimer. RSL oligomerization appears to be resistant to denaturation conditions. A smaller amount of dimers and monomers were also visible, probably because of partial oligomer denaturation (Supplementary Fig. 1B).

### Biophysical characterization of MOAβT and RSL-MOA by isothermal titration calorimetry (ITC)

MOAβT affinity towards Galα1-3Gal disaccharide, the terminal disaccharide of Galili epitope, was assayed by titration microcalorimetry. Large exothermic peaks were obtained at the beginning of the titration. The affinity was not strong enough to obtain a sigmoidal curve, and therefore, as recommended in such cases (Turnbull and Daranas 2003), the stoichiometry (N) was fixed, using a value of N = 2 since, as previously shown by Grahn et al., (2007), MOA binding sites have different affinities, and we expected at least two of them to be active. A dissociation constant (Kd) of 150 μM was obtained, which confirms the functionality of the isolated β-trefoil domain, and is in good agreement with the Kd measured for the whole MOA protein, i.e., 182 μM (Winter et al. 2002) (Fig. 2A).

**Figure 2:**
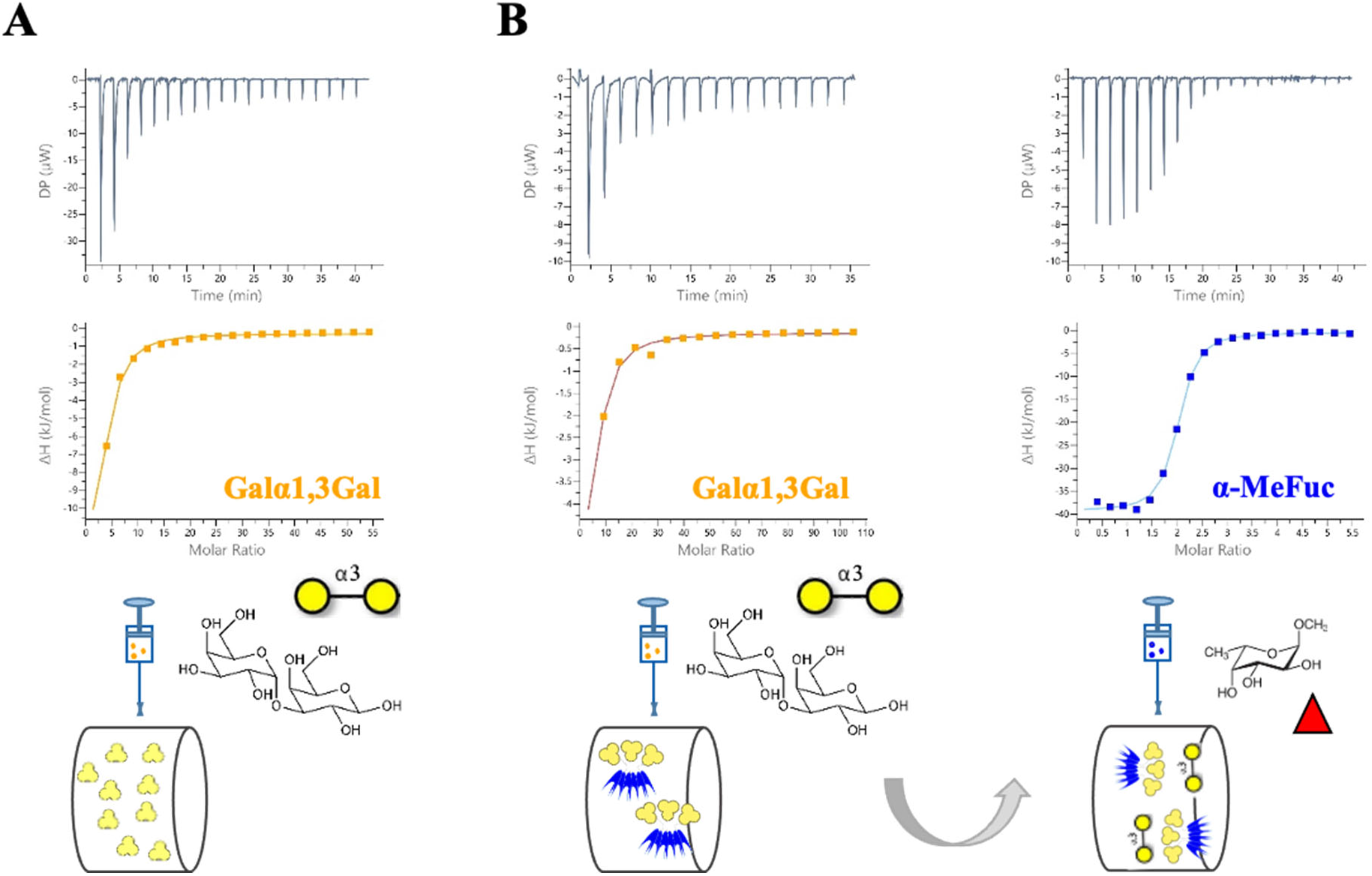
ITC thermographs of MOAβT and Janus lectin RSL-MOA. A) MOAβT binds Gala1-3Gal with a similar affinity as parental protein MOA. B) RSL-MOA can simultaneously bind two binding partners, i.e., Gala1-3Gal (yellow circles) and a-MeFuc (red triangle) confirming the activity of both functional domains.

RSL-MOA was designed to possess six binding sites for fucose and up to nine binding sites for α-galactose on the opposite faces. The functionality of both binding interfaces was tested by various biophysical approaches. To this end, we designed an ITC experiment with a consecutive injection of both ligands. First, αGal1-3Gal was titrated into the cell containing RSL-MOA. Subsequently, the cell content from the first experiment (complex RSL-MOA/Galα1-3Gal) was titrated by α-methyl fucoside (α-MeFuc) (Fig. 2B).

The thermogram obtained by titrating Galα1-3Gal in RSL-MOA solution was very similar to the one obtained for MOAβT. The dissociation constant (Kd = 226 μM) was comparable to the Kd for the isolated β-trefoil domain (above) and for the whole MOA (Winter et al. 2002). The subsequent titration by α-MeFuc resulted in a sigmoid shape, due to stronger affinity, with a measured stoichiometry of N= 2, corresponding to the presence of two fucose binding sites per RSL monomer. The Kd was measured to be 0.4 μM, in excellent agreement with the previously measured affinity (0.7 μM) for RSL (Kostlanová et al. 2005). This experiment proved that both parts of RSL-MOA are functional and able to bind their ligands at the same time.

### Biophysical characterization of RSL-MOA by surface plasmon resonance

To evaluate the effect of multivalency and therefore measure avidity instead of affinity, surface plasmon resonance (SPR) was used. Streptavidin CM5 chips were functionalized by different ligands, i.e., biotinylated PAA-α-fucose, PAA-α-galactose, and PAA-β-galactose.

No binding of RSL-MOA could be observed with α-galactose chip, even with high density surface(200 μg/mL biotinylated PAA-α-Gal), probably because of its very low affinity for the monosaccharide α-galactose (K_D_ = 8 mM) (Winter et al. 2002). On the opposite, RSL-MOA bound efficiently to fucosylated chips. In order to avoid mass transfer (de Mol and Fischer 2010), low density (LD) CM5 chip was prepared with mixture of PAA-α-fucose/PAA-β-galactose in the ratio 1:9 (200 μg/mL) and RSL-MOA was injected in increasing concentrations (5 to 1000 nM). The regeneration step was performed between each injection using 1 M fucose solution. As shown in Figure 3A, a dose-dependent response was obtained, with a steep association phase and very weak dissociation event. The steady-state analysis showed that even at the highest protein concentration (1000 nM) the chip surface was not saturated by RSL-MOA, i.e., the plateau phase was not reached (Fig. 3A). Nonetheless, the fitting of the kinetics constant could be performed and values for affinity (K_D_ = 4.9*10^-10^ M) and kinetics (k_on_ and k_off_) were estimated (Table 1).

**Figure 3:**
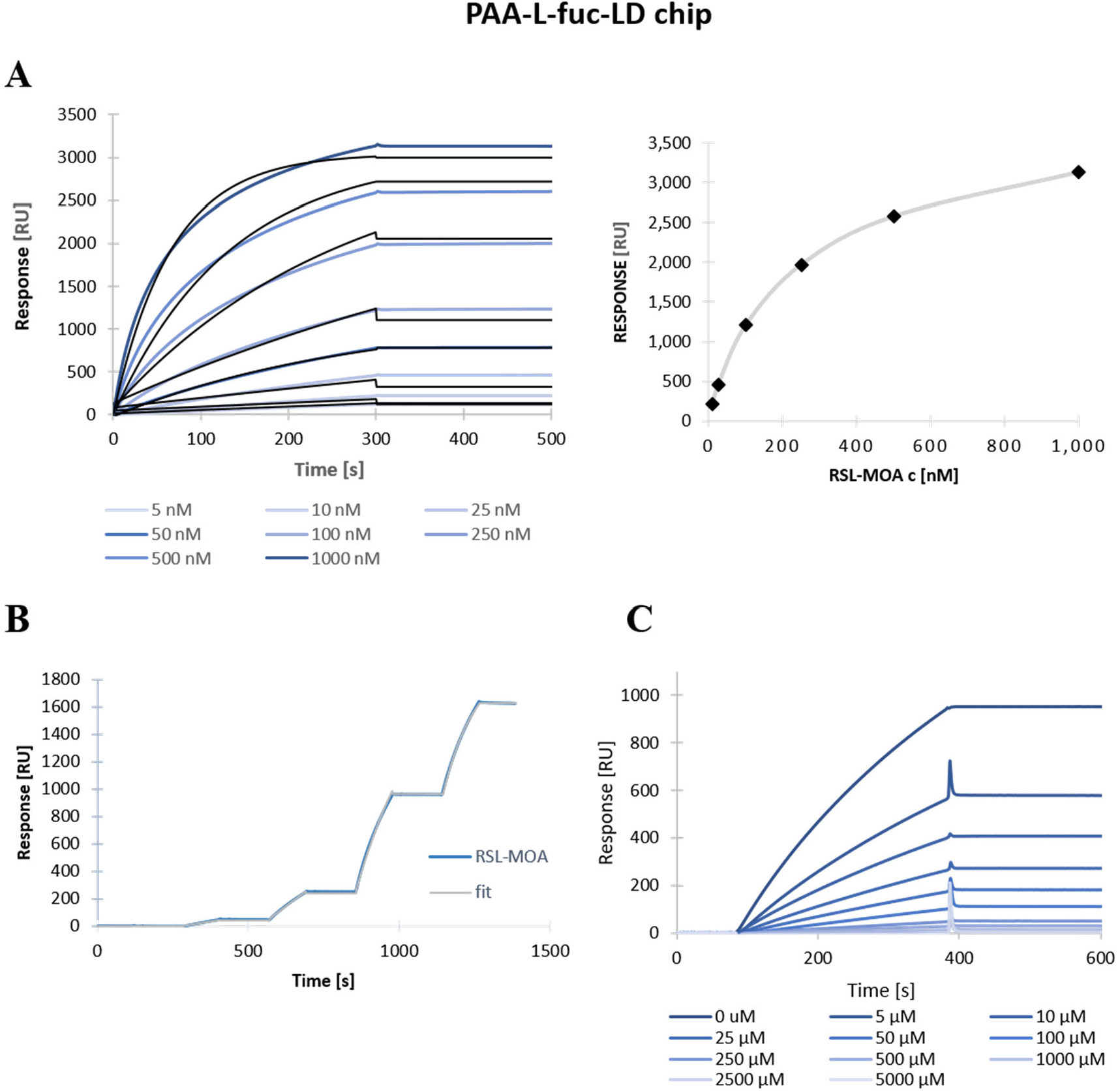
SPR sensorgrams of Janus lectin RSL-MOA on CM5-PAA-α-fuc LD chip. A) Titration experiment of different concentrations of RSL-MOA (blue) and fitting curves (1:1 binding fit) (black) (left) and steady-state analysis of RSL-MOA titration (right). B) Single-cycle kinetics of RSL-MOA with concentrations of 10, 50, 250, and 500 nM. C) Inhibition of 50 nM RSL-MOA by various concentration offucose.

**Table 1:** SPR statistics from titration and single-cycle kinetics experiments. The experiments were carried out on LD PAA-a-Fuc CM5 chips with a flow of 10 μL/min.

| Experimental condition | $k_{\text{on}}$<br>(1/Ms) | $k_{\text{off}}$<br>(1/s) | $K_D$<br>(M) | $R_{\text{MAX}}$<br>(RU) | $\text{Chi}^2$ |
| --- | --- | --- | --- | --- | --- |
| **Chip CM5 PAA-α-fuc LD**<br>*Classical kinetics* | $4.42 \cdot 10^2$ | $2.08 \cdot 10^{-7}$ | $4.9 \cdot 10^{-10}$ | 3030 | 8.23 |
| **Chip CM5 PAA-α-fuc LD**<br>*Single cycle kinetics* | $1.84 \cdot 10^4$ | $1.09 \cdot 10^{-7}$ | $5.9 \cdot 10^{-12}$ | 1960 | 40.4 |

As an alternative, a single-cycle kinetic experiment was performed. RSL-MOA was injected at four concentrations (10 to 500 nM) (Fig. 3B). The advantage of this method is that no regeneration step is needed. The estimated K_D_ of 5.9*10^-12^ M was two orders lower if compared to the titration experiment; however, such a variation is acceptable in binding events characterized by avidity. In order to evaluate the capacity of monosaccharides to inhibit the multivalent binding and compete with protein binding to the chip surface, 50 nM RSL-MOA was pre-incubated with various concentrations of fucose (5 to 5000 μM). As shown in Figure 3C, the complete inhibition was achieved in the presence of high concentrations of fucose (2500 and 5000 μM), and a IC50 value of 7.8 μM was obtained.

The ability of RSL-MOA to bind the glycan-decorated surface of SPR chips was confirmed by several experimental setups. The variations in the affinity constant just illustrate the difficulty to quantify binding when avidity is the major event. These results are in agreement with previous ones evaluating RSL avidity for the fucose chip (Arnaud et al. 2013). This avidity constant is at least 1000-fold higher than the affinity measured for fucose monosaccharide in solution, confirming the very strong cluster effect resulting from the topology of the RSL β-propeller with the presentation of six binding sites on the same face, binding very efficiently to fucose presented in a multivalent manner on a surface.

### MOAβT binds, while RSL-MOA binds and crosslinks differentially glyco-decorated giant unilamellar vesicles

The binding properties of MOAβT and RSL-MOA were tested with glyco-decorated giant unilamellar vesicles (GUVs). The GUVs were fluorescently labelled with Atto647N- or Atto488-DOPE lipid and contained functional spacer lipid (FSL) with different terminal saccharides attached, i.e. FSL-isoglobotriose, FSL-A, and FSL-B carrying Galα1-3Galβ1-4Glc, blood group A and blood group B terminal trisaccharides, respectively (Figure 4 A-D). These glyco-decorated GUVs were incubated with MOAβT and RSL-MOA lectins, which were both fluorescently labeled with same Atto488 fluorophore (Figure 4 A-D).

**Figure 4:**
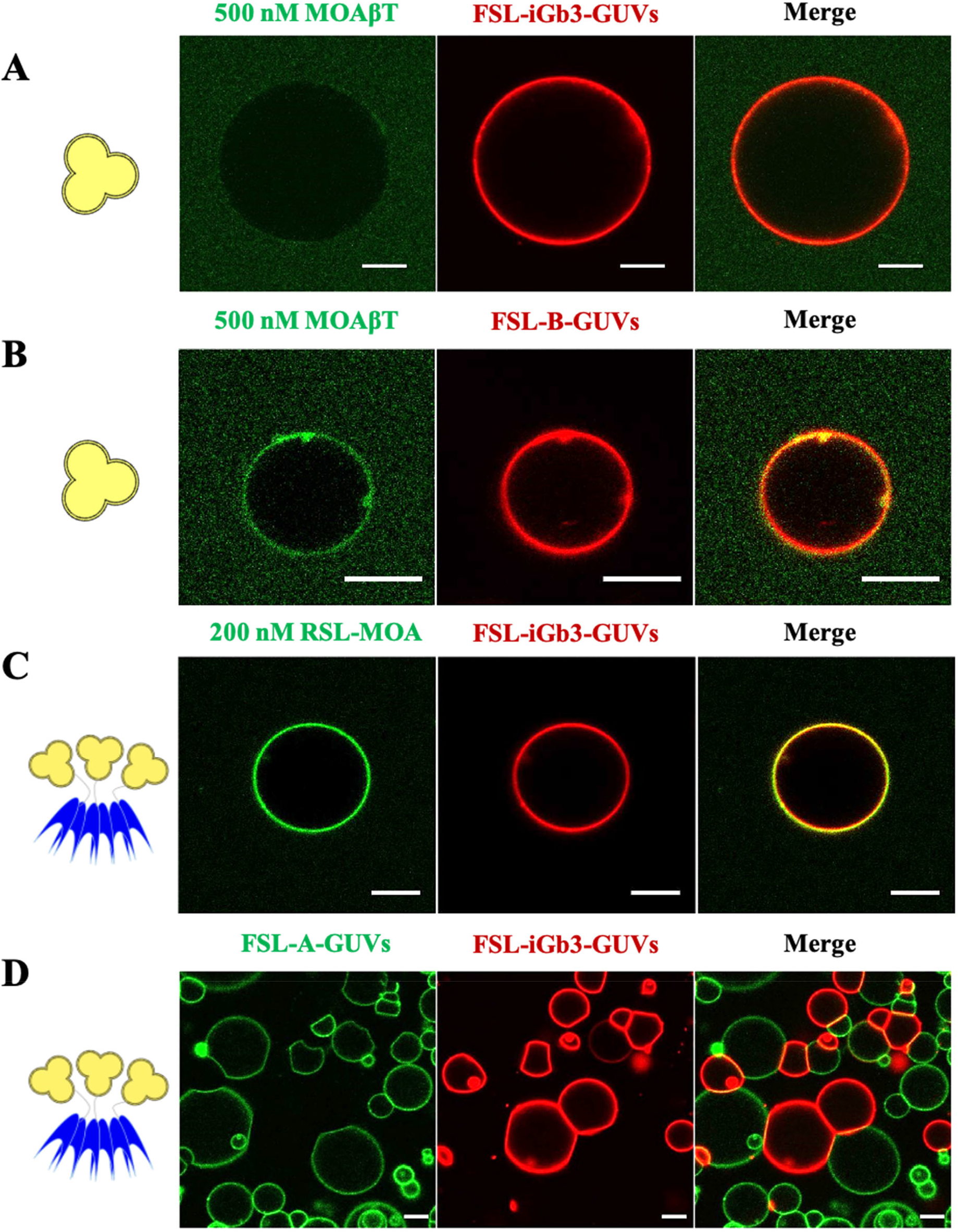
The binding properties of MOAβT and RSL-MOA to glyco-decorated GUVs. A) 500 nM MOAβT-Atto488 (green) shows almost no binding to the FSL-iGb3-GUVs (red, labelled with fluorescent lipid Atto647N-DOPE). B) 500 nM MOAβT-Atto488 (green) binds to FSL-B-GUVs (red, labelled with fluorescent lipid Atto647N-DOPE). C) 200 nM RSL-MOA-Atto488 (green) binds to FSL-iGb3-GUVs (red, labelled with fluorescent lipid Atto647N-DOPE). D) Unlabeled 200 nM RSL-MOA crosslinks two different populations GUVs FSL-iGb3-GUVs (red, labelled with fluorescent lipid Atto647N-DOPE) and FSL-A-GUVs (green, labelled with fluorescent lipid Atto488-DOPE) and crosslinks them. The GUVs were composed of DOPC, cholesterol, glycolipid of choice, and membrane dye to the proportion of 64.7:30:5:0.3 mol%, respectively. Scale bars are 10 μm.

MOAβT-Atto488 (500 nM, green) almost did not bind to FSL-iGb3-containing GUVs (red) (Fig. 4A). While, on the contrary, MOAβT-Atto488 bound FSL-B-containing GUVs and induced the tubule-like structures (Fig. 4B) indicating its ability to bind glycan structures on the membranes.

The Janus lectin RSL-MOA-Atto488 (green) binds efficiently to the surface of FSL-iGb3-GUVs (red), even at a concentration of 200 nM (Fig. 4C). The difference is striking when compared to the absence of binding of MOAβT-Atto488 (500 nM) (Fig. 4A
). The super-multivalency resulting from the presence of three β-trefoil domains, so in total nine possible αGal binding sites, makes a very strong difference in terms of ability to bind to the glycosylated surface.

The capacity of RSL-MOA to crosslink GUVs carrying different oligosaccharides was tested using two populations of fluorescently labeled vesicles, FSL-iGb3-GUVs (labelled with Atto647N-DOPE, red) and FSL-A-GUVs (labelled with Atto488-DOPE, green). When the unlabeled RSL-MOA was incubated with the liposomes, cross-linking is observed between red and green GUVs, as well as, cross-linking between GUVs of the same color. The cross-linking between same population GUVs is possibly due to lectin topology and multivalency, β-trefoil of MOA and β-propeller of RSL. Herein, due to the double specificity of lectin RSL-MOA, we confirmed that multivalent and two-site-oriented topology has clear potential in the crosslinking of glyco-decorated liposomes and proto-tissues (Fig. 4D).

### RSL-MOA binds to human epithelial cells

The ability of RSL-MOA to recognize and bind to glycans on the surface of human cells was investigated by flow cytometry. RSL-MOA was fluorescently labeled with a Cy5 dye (RSL-MOA-Cy5) and incubated with the non-small cell lung cancer cell line H1299, for 30 minutes at 4°C. As depicted in the histograms of fluorescence intensity (Fig. 5A), cells treated with increasing concentrations of RSL-MOA-Cy5 (0.07 – 0.7 μM) showed binding of the lectin to the cell surface in a dose-dependent manner. In order to confirm lectin specificity and glycan-driven binding to receptors at the plasma membrane of H1299 cells, a number of inhibition assays was performed. RSL-MOA-Cy5 was pre-incubated with 100 mM fucose or 100 mM synthetic analogue of α-Gal epitope, p-nitrophenyl-α-D-galactopyranoside (PNPG), respectively. The high concentrations of ligands were chosen due to previous observations with lectin RSL, for which a complete inhibition of binding to fucosylated receptors on the surface of H1299 cells was achieved only in the presence of 100 mM fucose (Siukstaite et al. 2021).

**Figure 5:**
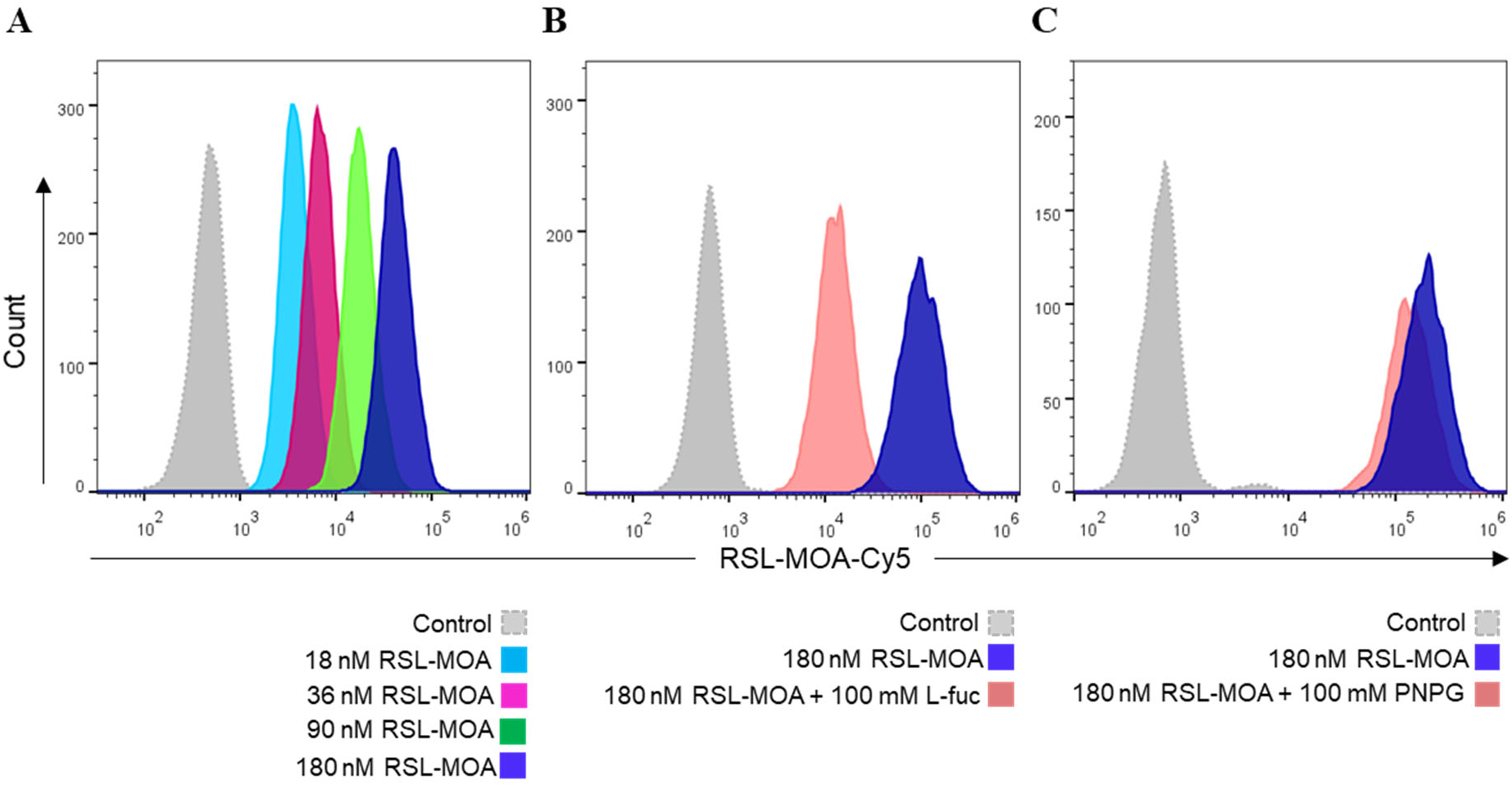
RSL-MOA shows dose-dependent binding to H1299 cells. A) Representative histogram plot of gated living H1299 cells pre-incubated with fluorescently labeled RSL-MOA-Cy5. Histograms of fluorescence intensity reveal a dose-dependent trend in binding according to used lectin concentrations. B) Representative histograms of fluorescence intensity of H1299 treated with RSL-MOA-Cy5 preincubated with 100 mM fucose. Inhibition of the RSL domain with high concentrations of fucose determines a shift in fluorescence intensity towards lower values (pink histogram), suggesting a reduced binding of lectin to H1299 cells. C) Representative histograms of fluorescence intensity of H1299 treated with RSL-MOA-Cy5 pre-incubated with 100 mM PNPG shows almost no reduction in lectin binding to cell surface. The control experiments correspond to samples of H1299 cells without any addition of lectin sample. The number of cells within the live population (y-axis) is plotted against the fluorescence intensity of RSL-MOA-Cy5 (x-axis).

The presence of 100 mM fucose significantly decreased RSL-MOA-Cy5 binding to treated cells by approx. 50% compared to unblocked RSL-MOA-Cy5 (Fig. 5B), but did not abolish it entirely. Increasing the concentration up to 500 mM of fucose did not result in a further decrease in protein binding. The interaction of the engineered lectin with the cell surface was therefore not completely inhibited upon saturation of the fucose-binding sites, suggesting that the MOA domain partially compensates for the binding of Janus lectin to H1299 cells. On the other hand, 100 mM PNPG had almost no effect on RSL-MOA-Cy5 binding to treated cells, as indicated by the histograms in Fig. 5C. Fluorescence of cells incubated with RSL-MOA-Cy5 or RSL-MOA-Cy5 pre-treated with PNPG exhibit a similar intensity, as illustrated by the overlapping histograms (Fig. 5C). Reasonably, inhibition of protein binding was not achieved due to the low affinity of MOA toward monosaccharides (Winter et al. 2002). These results suggest that both RSL and MOA domains of Janus lectin are able to bind to glycans present on H1299 cancer cells.

### Comparison of thermal stability of Janus lectins and their distinct domains

Janus lectins are created by assembling protein domains with different biophysical properties. It is therefore of interest to evaluate the effect of the fusion of these modules, and also to compare the biophysical properties of Janus lectins.

The thermal stability of RSL-MOA and its constituting domains was measured with differential scanning calorimetry (DSC). Thermograms of MOAβT and RSL are displayed in Figure 6. These two modules have very different thermal stability with a denaturation/midpoint temperature (Tm) of 42°C for the MOAβT and 92°C (with two Tm at 90.5°C and 93.8°C) for the β-propeller RSL. It could be that MOAβT is easily denatured as it represents only a domain of the whole original protein. The very strong thermal resistance of RSL is related to the compact and robust architecture of the β-propeller, and the two slightly different Tms extracted from the curve may correspond to the dissociation of the trimer followed by the unfolding of each monomer. As expected, adding α-MeFuc as a ligand to the protein solution increased the Tm of RSL to values of approximately 100°C and 103°C, for sugar concentrations of 125 and 250 μM, respectively.

**Figure 6:**
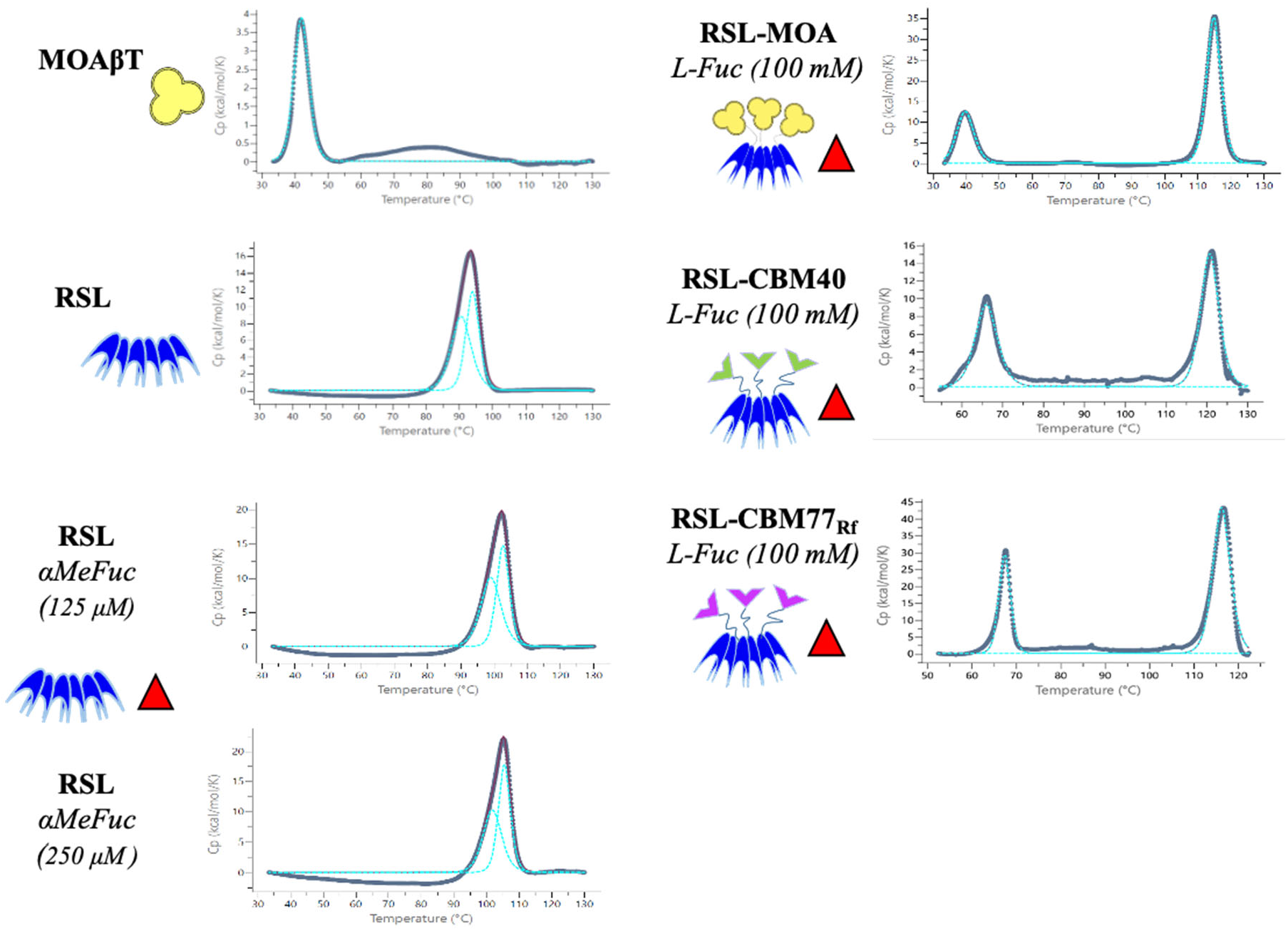
Thermal stability analysis of MOAβT, RSL, and Janus lectins RSL-MOA, RSL-CBM40, and RSL-CBM77_Rf_ with or without ligands. The figure displays experimental conditions, fitted data, and schematic representation of proteins and ligands.

The thermal unfolding of RSL-MOA and other Janus lectins was carried out in the presence of a high concentration of fucose since the protein was tested after the purification procedure. RSL-MOA displayed two events of denaturation at very different temperatures, i.e., 40°C and 115°C. From the results obtained on the separated domains, it is clear that the lower Tm corresponds to the unfolding of the MOA β-trefoil and the second event, with higher Tm, corresponds to the denaturation of RSL. The Tm of the MOA moiety is not different when isolated or linked to RSL. The Tm of the RSL moiety is higher than for the β-propeller alone, which is probably due to the high concentration of fucose in the medium.

Other Janus lectins were also assayed by DSC for comparison. The proteins RSL-CBM40 and RSL-CBM77Rf were obtained as described previously (Notova et al. 2022b; Ribeiro et al. 2018) and the buffer was supplemented by fucose as in the case of RSL-MOA. As already experienced with RSL-MOA, two separate events of denaturation were observed (Fig. 6) (Table 2). The high-temperature Tm, corresponding to the denaturation of RSL, is observed, with a value as high as 121°C for RSL-CBM40. CBM77_Rf_ and CBM40 have a similar β-sandwich fold, and they present rather similar melting temperatures of 67°C and 65°C, respectively. This value is more than 25 °C higher than the one observed for MOA, indicating a clear difference in thermal stability between β-sandwich and β-trefoil.

**Table 2:**
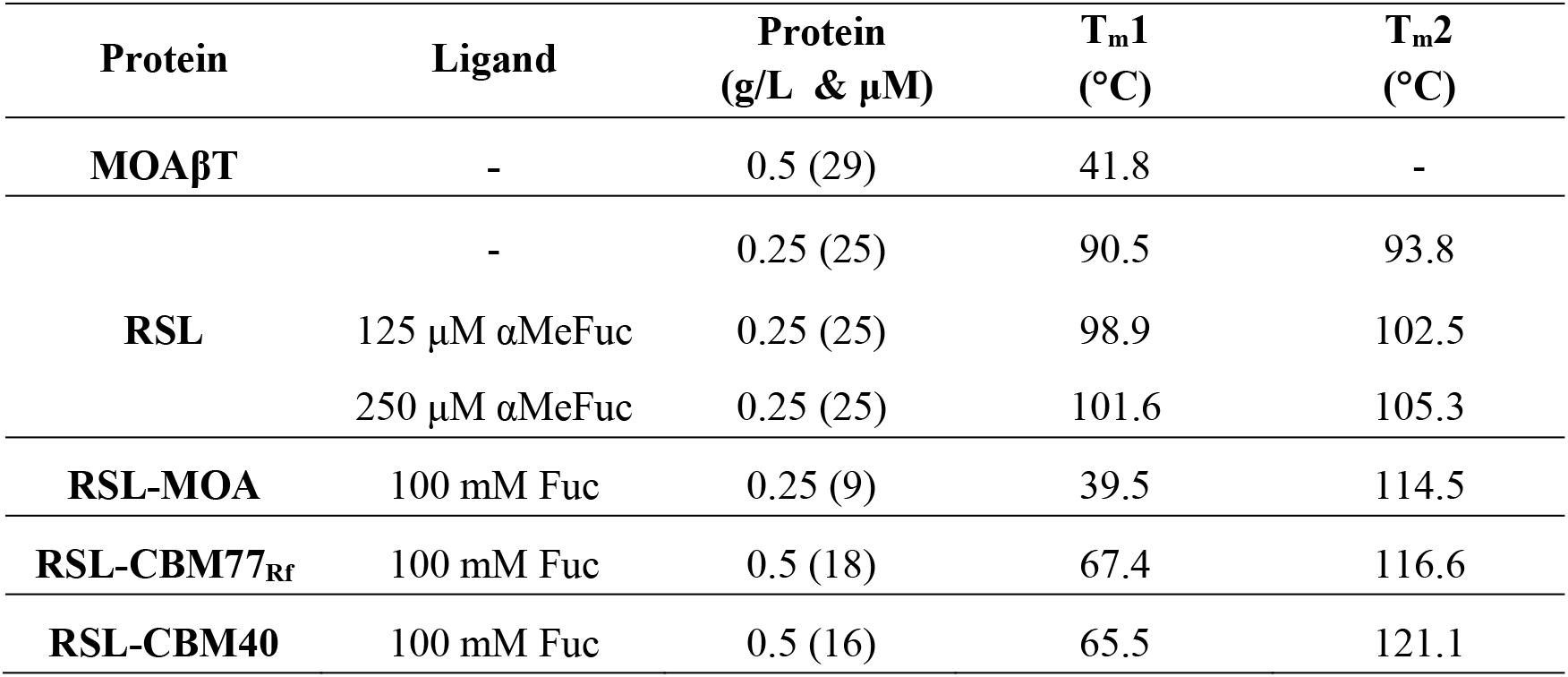
Thermal signatures of denaturation profiles of MOAβT, RSL, and Janus lectins. The scan rate was fixed at 200 °C/min.

## Discussion and Conclusion

In this work, we extended the concept of Janus lectin and therefore developed a universal strategy for increasing lectin valency and introducing an additional specificity (Notova et al. 2022b; Ribeiro et al. 2018). Janus lectins are engineered as fusion chimeras and due to the presence of the lectin RSL, these synthetic proteins assemble as trimers, resulting in the multiplication of lectin binding sites and thus expected higher affinity toward ligands. The first two Janus lectins, RSL-CBM40 (Ribeiro et al. 2018; Siukstaite et al. 2021) and RSL-CBM77Rf (Notova et al. 2022b) were designed as fusion chimeras of a lectin and CBM domain with a possible application as drug carriers or protein crosslinker in plant cell wall engineering, respectively. However, here we propose an alternative where the Janus lectin RSL-MOA is composed of two individual lectin domains, β-trefoil from lectin MOA and β-propeller of RSL.

The biophysical properties of RSL-MOA were compared with MOAβT, the engineered β-trefoil domain of MOA lectin. Both proteins showed the same affinity toward ligands in solution (ITC experiments) confirming their activity. However, their behavior was different when tested with glyco-decorated liposomes (GUV experiments). We observed that MOAβT showed almost no binding to FSL-iGb3-GUVs while RSL-MOA interacts with such vesicles and even crosslinks them. This is probably conditioned by the super-multivalency of engineered RSL-MOA that presents nine binding sites, instead of only three in MOAβT. On the other hand, MOAβT displayed active binding with blood group B GUVs (FSL-B-containing GUVs) confirming that the lectin domain is able to bind glycoconjugates on the membrane surface. RSL-MOA also has the ability to bind two different GUVs and crosslink them. This confirms that the engineered topology does not affect protein binding and such a lectin is of interest for the development of a highly selective tool for the construction of prototissues.

The SPR analysis of RSL-MOA revealed the strong dose-dependent binding of the β-propeller RSL toward low-density PAA-α-Fuc CM5 chips. Even though almost no dissociation event was observed, the fitting procedure allowed for the estimation of binding constants. Depending of the experimental design, some variations are observed in affinity and kinetics, due to the strong avidity of the system, generating complexity. Nonetheless, RSL-MOA showed at least 1000 higher affinity for surface-presented oligosaccharides compared to the solution state, supporting the fact that the addition of MOA does not affect its binding properties.

The ability of RSL-MOA to recognize glycans exposed at the surface of H1299 cells was analyzed by flow cytometry, demonstrating a strong dose-dependent binding. H1299 cells are characterized by the presence of highly fucosylated glycoconjugates, among others, on their surface (Jia et al. 2018). Furthermore, the effect of saturation of RSL-MOA sugar-binding sites with 100 mM fucose or 100 mM PNPG, was checked and resulted in a decrease in the interaction between lectin and cells. We believe that the presence of the galactose-specific MOA domain partially compensates the inhibition of fucose-dependent interaction between the lectin and the cell surface. Our hypothesis is supported by previous observations with the lectin RSL, which showed similar binding properties to H1299 cells that could be fully inhibited by 100 mM fucose. On the other hand, 100 mM PNPG did not affect RSL-MOA binding, suggesting that the synthetic analogue of α-galactose (PNPG) did not bind MOA with sufficient affinity, as already observed previously with α-galactose monosaccharide (Winter et al. 2002) and confirmed by SPR observations in the present work.

The thermal stabilities of three Janus lectins, i.e., RSL-MOA, RSL-CBM40, and RSL-CBM77_Rf_, were compared by DSC. During protein denaturation, two events of unfolding are observed for all Janus lectins implying the fact that each protein domain has a different stability. Structurally similar CBMs (β-sandwich fold) share almost the same Tm, while β-trefoil is much more thermally unstable. On the other hand, the β-propeller RSL showed extremely high Tm, whereas the addition of the ligand has a tremendous effect on its stability. Our findings, therefore, suggest that the structural fold of the individual domains should be taken into account when designing novel Janus lectins if stability is to be considered.

Until now, three Janus lectin with different specificities and architecture have been engineered. Therefore, we are confident that with this strategy, the novel bispecific lectins with improved valency can be obtained and find their applications in numerous fields of biotechnology and biomedicine.

## Methods

### Gene design and cloning

#### MOAβT

The gene for the β-trefoil domain of MOA was obtained by polymerase chain reaction where the plasmid pET-25b+RSL-MOA was used as a template. The primers, TTACATATGAGCTTACGTCGTGGC (Forward) and ATTACTCGAGTTACATGCGGTTGAAGTACC (Reverse) were designed to align to the full sequence of the β-trefoil domain of MOA and were ordered from Eurofins Genomics (Ebersberg, Germany). The restriction enzyme sites of NdeI and XhoI were added at 5’ and 3’ ends, respectively. Subsequently, the gene *moaβt* and plasmid pET-TEV vector (Houben et al. 2007) were digested by RE NdeI and XhoI and ligated resulting in the pET-TEV-MOAβT vector. After transformation by heat shock in the *E. coli* DH5α strain, a colony screening was performed, and the positive plasmids were amplified and controlled by sequencing.

#### RSL-MOA

The original amino acid sequence of the lectin MOA from *Marasmius oreades* was obtained from the PDB database. The gene *rsl-moa* was designed as a fusion chimera with RSL at the N-terminus and the β-trefoil domain of MOA at the C-terminus *via* the linker PNGELLSS. The gene was ordered from Eurofins Genomics (Ebersberg, Germany) after codon optimization for the expression in the bacteria *Escherichia coli*. The restriction enzyme sites of NdeI and XhoI were added at 5’ and 3’ ends, respectively. The synthesized gene was delivered in plasmid pEX-A2-RSL-MOA. Subsequently, plasmid pEX-A2-RSL-MOA and the pET-25b+ were digested by NdeI and XhoI restriction enzymes to ligate *rsl-moa* in pET-25b+. After transformation by heat shock in the *E. coli* DH5α strain, a colony screening was performed, and the positive plasmids were amplified and controlled by sequencing.

### Protein expression

*E. coli* BL21(DE3) cells were transformed by heat shock with pET-TEV-MOAβT plasmid prior pre-culture in Luria Broth (LB) media with 25 μg/mL kanamycin at 37°C under agitation at 180 rpm overnight. The next day, 10 mL of preculture was used to inoculate 1 L LB medium with 25 μg/mL kanamycin at 37°C and agitation at 180 rpm. When the culture reached OD_600nm_ of 0.6 - 0.8, the protein expression was induced by adding 0.1 mM isopropyl β-D-thiogalactoside (IPTG), and the cells were cultured at 16°C for 20 hours.

*E. coli* KRX (Promega) cells were transformed by heat shock with the pET-25b+-RSL-MOA plasmid and pre-cultured in LB media substituted with 50 μg/mL ampicillin at 37°C under agitation at 180 rpm overnight. The following day, 10 mL of preculture was used to inoculate 1 L LB medium with 50 μg/mL ampicillin at 37°C and agitation at 180 rpm. When reached an OD_600nm_ of 0.6 - 0.8, the protein expression was induced by adding 1% L-rhamnose, and the cells were cultured at 16°C for 20 hours.

The cells were harvested by centrifugation at 14000 × *g* for 20 min at 4°C and the cell paste was resuspended in 20 mM Tris/HCl pH 7.5, 100 mM NaCl (Buffer A), and lysed by a pressure cell disruptor (Constant Cell Disruption System) with a pressure of 1.9 kBar. The lysate was centrifuged at 24 000 × *g* for 30 min at 4°C and filtered on a 0.45 μm syringe filter prior to loading on an affinity column.

### Protein purification

#### MOAβT

The cell lysate was loaded on 1 mL HisTrap column (Cytiva) pre-equilibrated with Buffer A. The column was washed with Buffer A to remove all contaminants and unbound proteins. The MOA was eluted by Buffer A in steps during which the concentration of imidazole was increased from 25 mM to 500 mM. The fractions were analyzed by 12% SDS PAGE and those containing MOAβT were collected and deprived of imidazole by dialysis in Buffer A. The N-terminal His-tag was removed by TEV cleavage with the ratio 1:50 mg of TEV:protein in the presence of 0.5 mM EDTA and 1 mM TCEP over night at 19°C. After, the protein mixture was repurified on 1 mL HisTrap column (Cytiva) and the pure protein was concentrated by Pall centrifugal device with MWCO 3 kDa and stored at 4°C.

#### RSL-MOA

The cell lysate was loaded on 10 mL D-mannose-agarose resin (Merck) pre-equilibrated with Buffer A. The column was washed with Buffer A to remove all contaminants and unbound proteins and the flow-through was collected. RSL-MOA was eluted by Buffer A with the addition of 100 mM D-mannose or 100 mM L-fucose in one step. Due to not sufficient binding capacity of the column, the flow-through was reloaded on the column several times and the protein was eluted as described previously. The fractions were analyzed by 12% SDS PAGE and those containing RSL-MOA were collected and dialyzed against Buffer A. The protein was concentrated by Pall centrifugal device with MWCO 30 kDa and the pure protein fractions were pooled, concentrated, and stored at 4°C.

### Isothermal Titration Calorimetry (ITC)

ITC experiments were performed with MicroCaliTC200 (Malvern Panalytical). Experiments were carried out at 25°C ± 0.1°C. Protein and ligand samples were prepared in Buffer A. The ITC cell contained proteins in a concentration range from 0.05 mM to 0.2 mM. The syringe contained the ligand solutions in a concentration from 50 μM to 10 mM. 2 μL of ligands solutions were injected into the sample cell at intervals of 120 s while stirring at 750 rpm. Integrated heat effects were analyzed by nonlinear regression using one site binding model (MicroCal PEAQ-ITC Analysis software). The experimental data were fitted to a theoretical curve, which gave the dissociation constant (Kd) and the enthalpy of binding (ΔH).

### Surface plasmon resonance (SPR)

The SPR experiments were performed using a Biacore X100 biosensor instrument (GE Healthcare) at 25°C. Biotinylated polyacrylamide-attached (PAA) sugars, such as PAA-α-fucose, PAA-α-galactose, and PAA-β-galactose (Lectinity) were immobilized on CM5 chips (GE Healthcare) pre-coated with streptavidin, as previously described (Ribeiro et al., 2016). In the sample cell, the PAA-sugars were immobilized either as a mixture for low-density chips (1:9 of sugar of interest:non-interacting sugar) or as an individual PAA-sugar. The immobilization levels of chip CM5 LD PAA-α-fucose:PAA-β-galactose (1:9) were FC1: streptavidin 3694 RU, PAA-β-galactose 49 RU, FC2: streptavidin 3541 RU, PAA-α-fucose:PAA-β-galactose 596 RU. The solutions were prepared at a concentration 200 μg/mL in 10 mM HEPES buffer pH 7.5 with 100 mM NaCl and 0.05% Tween 20 (Buffer S). The reference cell was prepared in the same way as described for the sample cell, however, this time only non-interacting sugars were used. All the experiments were carried out in Buffer S. For the titration experiments, different concentrations of protein were injected onto the chip with the flow of 10 or 30 μL/min, and the protocol included steps: association time 300 s, dissociation time 300 s, 2x regeneration step for 180 s by 1 M fucose. The single-cycle kinetics experiment was carried out with the flow of 10 μL/min with an association and dissociation time of 120 s, while the regeneration was done only at the end of the run. For the inhibition assay, RSL-MOA with the concentration of 50 nM was pre-incubated with various concentrations of L-fucose for a minimum of 1 h at 4°C and subsequently injected onto the fucose chip. The used protocol was identical to the one used for titration experiments. The data analysis was performed by BIAevaluation software.

### Protein labeling

RSL-MOA and MOAβT were dissolved at 1 mg/mL in Dulbecco’s phosphate-buffered saline (PBS) and stored at 4°C prior to usage. For fluorescent labeling, NHS-ester conjugated Atto488 (Thermo Fisher) or Cy5 (GE Healthcare) were used. Fluorescent dyes were dissolved at a final concentration of 10 mg/mL in water-free DMSO (Carl RothGmbH & Co), aliquoted, and stored at −20 °C before usage according to the manufacturer’s protocol. For the labeling reaction, 100 μL of lectin (1 mg/mL) was supplemented with 10 μL of a 1 M NaHCO_3_ (pH 9.0) solution. Hereby, the molar ratio between dye and lectin was 5:1 for RSL-MOA (cell assays) and 2:1 for RSL-MOA and MOAβT for GUV assays. The labeling mixture was incubated at 4°C for 90 min, and uncoupled dyes were separated using Zeba Spin™ desalting columns (7 kDa MWCO, 0.5 mL, Thermo Fischer). Labeled lectins were stored at 4°C, protected from light.

### Composition and preparation of giant unilamellar vesicles (GUVs)

GUVs were composed of 1,2-dioleoyl-sn-glycero-3-phosphocholine (DOPC), cholesterol (both AvantiPolar Lipids, United States), Atto647N 1,2-dioleoyl-sn-glycero-3-phosphoethanolamine (DOPE; Sigma Aldrich), Atto488 1,2-dioleoyl-sn-glycero-3-phosphoethanolamine (DOPE; Sigma Aldrich) and one of the following glycolipids at a molar ratio of 64.7:30:0.3:5. The glycolipids are FSL-A (Function-Spacer-Lipid with blood group A trisaccharide) (SigmaAldrich), FSL-B (Function-Spacer-Lipid with blood group B trisaccharide) (Sigma Aldrich) or FSL-isoGb3 (Function-Spacer-Lipid with iso-globotriaosyl saccharide).

GUVs were prepared by the electroformation method as earlier described (Madl et al., 2016). Briefly, lipids dissolved in chloroform of a total concentration of 0.5 mg/mL were spread on indium tin oxid-covered (ITO) glass slides and dried in a vacuum for at least one hour or overnight. Two ITO slides were assembled to create a chamber filled with sucrose solution adapted to the osmolarity of the imaging buffer of choice, either HBSS (for live-cell imaging) or PBS (for GUV-only imaging). Then, an alternating electrical field with a field strength of 1 V/mm was implemented for 2.5 hours at RT. Later we observed the GUVs in chambers manually built as described (Madl et al., 2016).

### Imaging of RSL-MOA and MOAβT binding to GUVs

Samples containing GUVs and lectins were imaged using a confocal fluorescence microscope (Nikon Eclipse Ti-E inverted microscope equipped with Nikon A1R confocal laser scanning system, 60x oil immersion objective, NA = 1.49, and four laser lines: 405 nm, 488 nm, 561 nm, and 640 nm). Image acquisition and processing were made using the software NIS-Elements (version 4.5, Nikon) and open-source Fiji software (https://imagej.net/software/fiji/).

### Cell culture

The human lung epithelial cell line H1299 (American Type Culture Collection, CRL-5803) was cultured in Roswell Park Memorial Institute (RPMI) medium supplemented with 10% fetal calf serum (FCS) and 4 mM L-glutamine at 37°C and 5% CO2, under sterile conditions. Cells were cultivated in standard TC-dishes 100 (Sarstedt AG & Co. KG) until 90% confluence, detached with trypsin (0.05% trypsin-EDTA solution; Sigma-Aldrich Chemie GmbH) and reseeded for a subculture or for experiments. For experiments, cells were incubated with different concentrations of RSL-MOA for indicated time points.

### Flow Cytometry Analysis

H1299 cells were detached with 2 mL of 1.5 mM EDTA in PBS, and 1 × 10^5^ cells were counted and transferred to a U-bottom 96 well plate (Sarstedt AG & Co. KG). To quantify protein binding to cell surface receptors, cells were incubated with different concentrations of fluorescently labeled RSL-MOA-Cy5 lectin for 30 min at 4°C and protected from light compared to PBS-treated cells as a negative control. For the saturation of glycan-binding sites, 180 nM RSL-MOA-Cy5 was preincubated with 100 mM soluble L-fucose or with 100 mM 4-Nitrophenyl α-D-galactopyranoside (PNPG; Sigma-Aldrich, Chemie GmbH), for 30 min at RT and in the absence of light. At the end of pre-incubation, the solution was diluted 100 times and added to cells for 30 min at 4°C, in the dark. Subsequently, cells were centrifuged at 1600 × g for 3 min at 4°C and washed twice with FACS buffer (PBS supplemented with 3% FCS v/v). After the last washing step, the cells were resuspended with FACS buffer and transferred to FACS tubes (Kisker Biotech GmbH Co. KG) on ice and protected from light. The fluorescence intensity of treated cells was monitored at FACS Gallios (Beckman Coulter Inc.) and further analyzed using FlowJo V.10.5.3.

### Differential scanning calorimetry (DSC)

The DCS experiments were carried out in Micro-Cal PEAQ DSC instrument (Malvern Panalytical). The protein samples were prepared in Buffer A, which could be substituted with the addition of a ligand. The sample cell contained the protein solutions in the concentration range of 9-29 mM. The reference cell contained the same buffer as present in the sample but without protein. The increase in temperature was measured from 20-130°C with a scan rate of 200°C/min. The data were analyzed by Micro-Cal PEAQ DSC software with a non-two-state and progress baseline method. fitting model.

## Acknowledgment

This research was funded by the European Union Horizon 2020 Research and Innovation Program under the Marie Skłodowska-Curie grant agreement synBIOcarb (No. 814029). AI acknowledges support from Glyco@Alps (ANR-15-IDEX-02) and Labex Arcane/CBH-EUR-GS (ANR-17-EURE-0003). Moreover, WR acknowledges support by the Ministry for Science, Research and Arts of the State of Baden-Württemberg (Az: 33-7532.20), and by the Freiburg Institute for Advanced Studies (FRIAS). This publication is partially based upon work from COST Action CA18103 (INNOGLY), supported by COST (European Cooperation in Science and Technology).

## Bibliography

Arnaud J, Claudinon J, Tröndle K, Trovaslet M, Larson G, Thomas A, Varrot A, Römer W, Imberty A, Audfray A. 2013. Reduction of lectin valency drastically changes glycolipid dynamics in membranes but not surface avidity. ACS Chem. Biol., 8:1918–1924, 10.1021/cb400254b

Bonnardel F, Mariethoz J, Salentin S, Robin X, Schroeder M, Pérez S, Lisacek F, Imberty A. 2019. UniLectin3D, a database of carbohydrate binding proteins with curated information on 3D structures and interacting ligands. Nucleic Acids Res., 47:D1236–D1244, 10.1093/nar/gky832

Chung CH, Mirakhur B, Chan E, Le QT, Berlin J, Morse M, Murphy BA, Satinover SM, Hosen J, Mauro D, et al. 2008. Cetuximab-induced anaphylaxis and IgE specific for galactose-alpha-1,3-galactose. N. Engl. J. Med., 358:1109–1117, 10.1056/NEJMoa074943

Cooper DK, Koren E, Oriol R. 1994. Oligosaccharides and discordant xenotransplantation. Immunol. Rev., 141:31–58, 10.1111/j.1600-065x.1994.tb00871.x

Cordara G, Van Eerde A, Grahn EM, Winter HC, Goldstein IJ, Krengel U. 2016. An unusual member of the papain superfamily: Mapping the catalytic cleft of the Marasmius oreades agglutinin (MOA) with a caspase inhibitor. PLoS ONE, 10.1371/journal.pone.0149407

de Mol NJ, Fischer MJ. 2010. Surface plasmon resonance: a general introduction. Methods Mol. Biol., 627:1–14, 10.1007/978-1-60761-670-2_1

Estola E, Elo J. 1952. Occurrence of an exceedingly weak A blood group property in a family. Ann. Med. Exp. Biol. Fenn., 30:79–87,

Fettis MM, Farhadi SA, Hudalla GA. 2019. A chimeric, multivalent assembly of galectin-1 and galectin-3 with enhanced extracellular activity. Biomater Sci, 7:1852–1862, 10.1039/c8bm01631c

Galili U, Shohet SB, Kobrin E, Stults CL, Macher BA. 1988. Man, apes, and Old World monkeys differ from other mammals in the expression of alpha-galactosyl epitopes on nucleated cells. J. Biol. Chem., 263:17755–17762

Grahn E, Askarieh G, Holmner A, Tateno H, Winter HC, Goldstein IJ, Krengel U. 2007. Crystal structure of the Marasmius oreades mushroom lectin in complex with a xenotransplantation epitope. J. Mol. Biol., 369:710–721, 10.1016/j.jmb.2007.03.016

Hazes B. 1996. The (QxW)3 domain: a flexible lectin scaffold. Protein Sci., 5:1490–1501, 10.1002/pro.5560050805

Houben K, Marion D, Tarbouriech N, Ruigrok RW, Blanchard L. 2007. Interaction of the C-terminal domains of sendai virus N and P proteins: comparison of polymerase-nucleocapsid interactions within the paramyxovirus family. J. Virol., 81:6807–6816, 10.1128/JVI.00338-07

Irumagawa S, Hiemori K, Saito S, Tateno H, Arai R. 2022. Self-Assembling Lectin Nano-Block Oligomers Enhance Binding Avidity to Glycans. Int J Mol Sci, 23, 10.3390/ijms23020676

Jia L, Zhang J, Ma T, Guo Y, Yu Y, Cui J. 2018. The Function of Fucosylation in Progression of Lung Cancer. Front Oncol, 8:565, 10.3389/fonc.2018.00565

Juillot S, Cott C, Madl J, Claudinon J, Van Der Velden NSJ, Künzler M, Thuenauer R, Römer W. 2016. Uptake of Marasmius oreades agglutinin disrupts integrin-dependent cell adhesion. Biochim. Biophys. Acta, 1860:392–401, 10.1016/j.bbagen.2015.11.002

Kostlanová N, Mitchell EP, Lortat-Jacob H, Oscarson S, Lahmann M, Gilboa-Garber N, Chambat G, Wimmerová M, Imberty A. 2005. The fucose-binding lectin from Ralstonia solanacearum: a new type of b-propeller architecture formed by oligomerisation and interacting with fucoside, fucosyllactose and plant xyloglucan. J. Biol. Chem., 280:27839–27849, 10.1074/jbc.M505184200

Kruger RP, Winter HC, Simonson-Leff N, Stuckey JA, Goldstein IJ, Dixon JE. 2002. Cloning, expression, and characterization of the Galalpha 1,3Gal high affinity lectin from the mushroom Marasmius oreades. J. Biol. Chem., 277:15002–15005, 10.1074/jbc.M200165200

Macher BA, Galili U. 2008. The Galalpha1,3Galbeta1,4GlcNAc-R (alpha-Gal) epitope: a carbohydrate of unique evolution and clinical relevance. Biochim. Biophys. Acta, 1780:75–88, 10.1016/j.bbagen.2007.11.003

Murzin AG, Lesk AM, Chothia C. 1992. beta-Trefoil fold. Patterns of structure and sequence in the Kunitz inhibitors interleukins-1 beta and 1 alpha and fibroblast growth factors. J. Mol. Biol., 223:531–543, 10.1016/0022-2836(92)90668-a

Notova S, Bonnardel F, Lisacek F, Varrot A, Imberty A. 2020. Structure and engineering of tandem repeat lectins. Curr. Opin. Struct. Biol., 62:39–47, 10.1016/j.sbi.2019.11.006

Notova S, Bonnardel F, Rosato F, Siukstaite L, Schwaiger J, Bovin N, Varrot A, Römer W, Lisacek F, Imberty aA. 2022a. A pore-forming β-trefoil lectin with specificity for the tumor-related glycosphingolipid Gb3. BioRXiv, 10.1101/2022.02.10.479907

Notova S, Cannac N, Rabagliati L, Touzard M, Mante J, Navon Y, Coche-Guérente L, Lerouxel O, Heux L, Imberty A. 2022b. Building artificial plant cell wall on lipid bilayer by assembling polysaccharides and engineered proteins. BioRXiv, 10.1101/2022.07.25.501355

Oh YJ, Dent MW, Freels AR, Zhou Q, Lebrilla CB, Merchant ML, Matoba N. 2022. Antitumor activity of a lectibody targeting cancer-associated high-mannose glycans. Mol Ther, 30:1523–1535, 10.1016/j.ymthe.2022.01.030

Ramberg KO, Guagnini F, Engilberge S, Wronska MA, Rennie ML, Perez J, Crowley PB. 2021. Segregated Protein-Cucurbit[7]uril Crystalline Architectures via Modulatory Peptide Tectons. Chemistry, 27:14619–14627, 10.1002/chem.202103025

Ribeiro JP, Villringer S, Goyard D, Coche-Guerente L, Höferlin M, Renaudet O, Römer W, Imberty A. 2018. Tailor-made Janus lectin with dual avidity assembles glycoconjugate multilayers and crosslinks protocells. Chem. Sci., 9:7634–7641 10.1039/C8SC02730G

Ross JF, Wildsmith GC, Johnson M, Hurdiss DL, Hollingsworth K, Thompson RF, Mosayebi M, Trinh CH, Paci E, Pearson AR, et al. 2019. Directed Assembly of Homopentameric Cholera Toxin B-Subunit Proteins into Higher-Order Structures Using Coiled-Coil Appendages. J. Am. Chem. Soc., 141:5211–5219, 10.1021/jacs.8b11480

Sharon N, Lis H. 2004. History of lectins: from hemagglutinins to biological recognition molecules. Glycobiology, 14:53R–62R, 10.1093/glycob/cwh122

Siukstaite L, Rosato F, Mitrovic A, Müller PF, Kraus K, Notova S, Imberty A, Römer W. 2021. The two sweet sides of Janus lectin drive crosslinking of liposomes to cancer cells and material uptake Toxins, 13:792, 10.3390/toxins13110792

Tateno H, Goldstein IJ. 2004. Partial identification of carbohydrate-binding sites of a Galalpha1,3Galbeta1,4GlcNAc-specific lectin from the mushroom Marasmius oreades by site-directed mutagenesis. Arch. Biochem. Biophys., 427:101–109, 10.1016/j.abb.2004.04.013

Terada D, Kawai F, Noguchi H, Unzai S, Hasan I, Fujii Y, Park SY, Ozeki Y, Tame JRH. 2016. Crystal structure of MytiLec, a galactose-binding lectin from the mussel Mytilus galloprovincialis with cytotoxicity against certain cancer cell types. Sci. Rep., 6:28344, 10.1038/srep28344

Turnbull WB, Daranas AH. 2003. On the value of c: can low affinity systems be studied by isothermal titration calorimetry? J. Am. Chem. Soc., 125:14859–14866, 10.1021/ja036166s

Ward EM, Kizer ME, Imperiali B. 2021. Strategies and Tactics for the Development of Selective Glycan-Binding Proteins. ACS Chem. Biol., 16:1795–1813, 10.1021/acschembio.0c00880

Winter HC, Mostafapour K, Goldstein IJ. 2002. The mushroom Marasmius oreades lectin is a blood group type B agglutinin that recognizes the Galalpha 1,3Gal and Galalpha 1,3Galbeta 1,4GlcNAc porcine xenotransplantation epitopes with high affinity. J. Biol. Chem., 277:14996–15001, 10.1074/jbc.M200161200

