## Supplementary material for "Extending Janus lectins architecture: characterization and application to protocells": Supp Figure 1

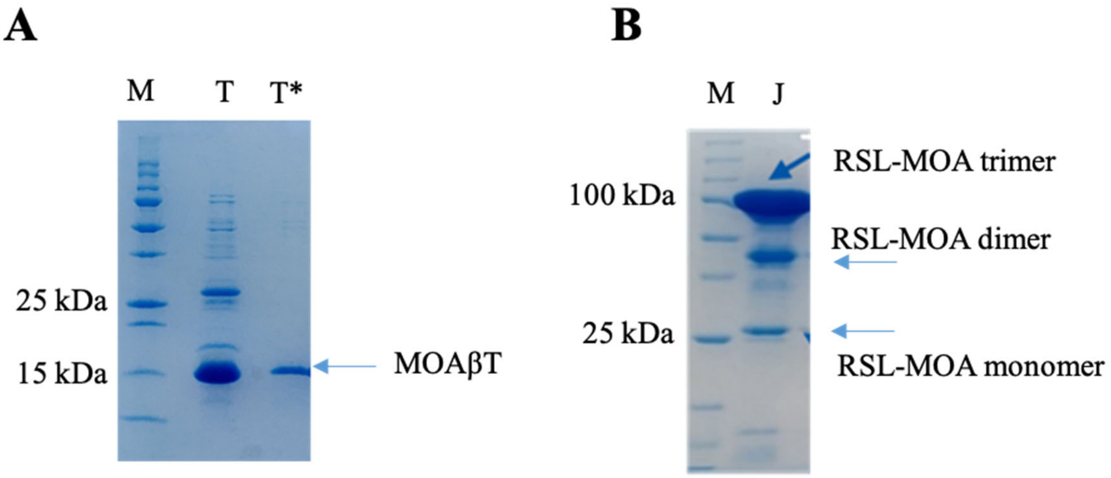


***Supplementary figure 1: SDS PAGE analysis of MOAβT and RSL-MOA.*** *The protein samples were analyzed under denaturing conditions on 12% polyacrylamide gel. M-protein marker, J-Janus lectin, T- MOAβT, T*- MOAβT cleaved by TEV protease. A) MOAβT with the estimated size of 17.2 kDa appears as monomeric and the impurities were eliminated during TEV cleavage and subsequent purification of protein. B) Janus lectin RSL-MOA is resistant to denaturation conditions and on the gel appears as a monomer (28 kDa), dimer (56 kDa), and trimer (83 kDa) whereas the trimer is the most abundant.*
